# Innovative CRISPR Etf-QO Mutant Knock-in Model Mimics MADD-associated Lipid Storage Myopathy and Reveal Therapeutic Potential of Exercise Intervention

**DOI:** 10.64898/2026.05.18.726022

**Authors:** Sachin Budhathoki, Yiming Guo, Mary Doamekpor, Girish Melkani

## Abstract

Multiple acyl-CoA dehydrogenase deficiency (MADD) is a rare lipid storage myopathy caused predominantly by pathogenic variants in ETFDH, which encodes ETF-QO in *Drosophila*, a mitochondrial electron-transfer protein required for fatty acid β-oxidation. Impaired ETF-QO function disrupts electron transfer from multiple acyl-CoA dehydrogenases to the ubiquinone pool, resulting in lipid accumulation, mitochondrial dysfunction, and progressive neuromuscular and cardiac abnormalities. Despite the availability of animal models of fatty acid oxidation disorders, genetically precise in vivo models carrying patient-relevant ETFDH missense variants and permitting longitudinal assessment of disease progression remain limited. Here, we generated CRISPR/Cas9 knock-in *Drosophila melanogaster* models of late-onset MADD harboring three conserved Etf-QO missense substitutions (*L138R*, *S307C*, and *L409F*), corresponding to human ETFDH mutations *L127R*, *S296C*, and *L399F*, respectively. These variants are located to the conserved FAD- and ubiquinone-binding regions of ETF-QO. Etf-QO mutant flies developed progressive deficits in locomotor activity and skeletal muscle performance accompanied by marked lipid droplet accumulation in indirect flight muscles, cardiac tissue, and fat bodies. Cardiac phenotyping further revealed impaired function, characterized by reduced fractional shortening, prolonged heart period, and increased arrhythmicity index. Consistent with mitochondrial bioenergetic dysfunction, Etf-QO mutants exhibited reduced oxygen consumption, increased oxidative stress, and decreased ATP levels. Molecular analyses indicated activation of cellular energy- and mitochondrial-stress responses, including increased AMPK and PGC-1α signaling, together with elevated Pink1, Parkin, and SOD2 expression, suggesting engagement of mitochondrial quality-control and antioxidant defense pathways in response to ETF-QO dysfunction. Importantly, exercise paradigm consisting of 15 min of daily exercise for 2.5 weeks significantly improved locomotor, skeletal muscle, and cardiac performance while reducing lipid accumulation and reactive oxygen species burden. Together, these findings establish innovative CRISPR knock-in *Drosophila* models that recapitulate key neuromuscular, metabolic, mitochondrial, and cardiac features of ETFDH-associated MADD. The models reveal coordinated mitochondrial stress and energy-signaling responses to ETF-QO dysfunction and demonstrate that exercise can ameliorate multiple disease phenotypes. These findings provide a tractable *in vivo* platform for dissecting MADD pathogenesis and evaluating potential therapeutic strategies targeting mitochondrial dysfunction and lipid storage myopathies.

## Introduction

Lipid Storage Myopathies (LSMs) are a heterogeneous group of inherited metabolic disorders characterized by abnormal accumulation of lipids within muscle cells, leading to progressive skeletal muscle weakness and cardiomyopathy^1–3^. These disorders arise primarily from disruptions in mitochondrial fatty acid β-oxidation (FAO), lipid trafficking, or electron transfer processes required for oxidative metabolism^4, 5^. Impairments in lipid trafficking and electron transfer affect the use of fatty acids as an energy source, resulting in ectopic lipid accumulation, reduced Adenosine Triphosphate (ATP) production, and progressive dysfunction of high-energy tissues such as skeletal muscle and heart^6^. Clinically, LSM spectrum ranges from fatal neonatal and congenital forms to progressive adult-onset types^7^. Among LSMs, MADD, also known as Glutaric Acidemia Type II, represents a paradigmatic disorder that links impaired mitochondrial FAO to systemic metabolic dysfunction^8^. MADD is caused by pathogenic variants in genes encoding electron transfer flavoprotein subunit A (ETFA), electron transfer flavoprotein subunit B (ETFB), or most commonly ETFDH^9^. ETFDH functions as a key mitochondrial enzyme that transfers electrons from multiple acyl-CoA dehydrogenases to the ubiquinone pool of the respiratory chain^10^. Disruption of this step compromises FAO across short-, medium-, and long-chain substrates, leading to accumulation of lipid intermediates, impaired oxidative phosphorylation, and cellular energy stress^11^. Mutations in ETFA and ETFB are mostly associated with severe neonatal-onset form characterized by dysfunction in mitochondrial energy metabolism and early mortality^12^. In contrast, mutations in ETFDH often result in late-onset MADD with partially retained enzymatic activity^8^. Late-onset MADD patients commonly present with progressive proximal muscle weakness, recurrent rhabdomyolysis, myalgia, and exercise intolerance and cardiomyopathy in some case-reported study^13–16^. Because FAO is a dominant energy source during prolonged exercise, defects in electron transfer exacerbate metabolic stress and increase susceptibility to oxidative damage^17^. Cardiac tissues subjected to oxidative stress exhibit increased ROS, DNA-damage signaling, and impaired structural integrity^18^. Although exercise intolerance is common in late-onset MADD, available clinical observations indicate that it reflects reduced metabolic flexibility rather than a fixed inability to perform physical activity^19^. Many patients recover exercise capacity once metabolic stability is achieved, particularly those with riboflavin-responsive variants^20^. Physical activity activates AMP-activated protein kinase (AMPK), a key energy sensor that stimulates peroxisome proliferator-activated receptor gamma coactivator 1-alpha (PGC-1α)^21, 22^. However, the molecular and physiological consequences of exercise in the context of ETFDH dysfunction remain poorly defined.

Current therapeutic options for MADD are limited and largely supportive. Riboflavin (flavin) supplementation represents the most effective treatment for a subset of patients, particularly those harboring flavin-responsive ETFDH missense mutations^23^. Riboflavin is thought to enhance FAD binding, stabilize mutant ETFDH protein, and partially restore electron transfer function^24^. Nevertheless, not all late-onset mutations respond to riboflavin, and many patients continue to experience progressive muscle weakness and cardiomyopathy despite treatment. Dietary fat restriction, carnitine supplementation, and coenzyme Q10 have shown variable benefit, underscoring the need for improved mechanistic understanding and complementary therapeutic strategies^25^. Progress in understanding MADD pathogenesis has been limited by the lack of genetically precise and physiologically scalable *in vivo* models that capture patient-specific mutations and age-dependent disease progression. Models incorporating patient-relevant ETFDH missense mutations remain scarce, as most existing systems rely on acute knockdown, non-physiological overexpression, or pharmacological inhibition which cannot capture the genotype-phenotype relationships characteristic of late-onset MADD. In this context, genetically tractable systems that preserve endogenous regulation while permitting longitudinal analysis are critically needed.

*Drosophila melanogaster* (commonly referred to as fruit-fly) has emerged as a powerful model organism for studying metabolic diseases affecting skeletal muscle and heart, owing to its high conservation of mitochondrial pathways, well-characterized striated muscle and cardiac physiology^26–31^. Importantly, the fly ortholog of ETFDH, ETF-ubiquinone oxidoreductase (Etf-QO), retains conserved FAD- and ubiquinone-binding domains and mutations that are disrupted by MADD-associated mutations^10, 32^. In this study, we generated the first CRISPR knock-in *Drosophila* models carrying clinically relevant missense mutations in Etf-QO, the conserved ortholog of human ETFDH, to model late-onset MADD under endogenous regulation. Using locomotor and muscle performance assays, lipid imaging, metabolic and gene expression analyses, and high-resolution cardiac physiology, we perform phenotypic and molecular characterization of the mutation. We further assess impact of moderate exercise as a non-pharmacological modifier and show that it improves skeletal muscle and cardiac performance while reducing lipid accumulation and oxidative stress, supporting adaptive mitochondrial remodeling as a key response to ETFDH dysfunction. Overall, these findings identify conserved disease phenotypes and mechanisms in LSM and indicate that structured endurance exercise produces measurable functional benefits under Etf-QO mutation backgrounds.

## Materials and Methods

### 1. Drosophila Stocks and Standard Environmental Conditions

CRISPR knock-in *Drosophila* models were developed to explore the effects of mutations in the Etf-QO gene associated with MADD. The specific point mutations introduced were Etf-QO^L127R^, Etf-QO^S296C^, and Etf-QO^L399F^ associated with FAD and ubiquinone binding domains, recapitulating mutations observed in human cases of late-onset MADD. The point mutations were generated with the help of Rainbow Transgenic Flies, Inc. through microinjection of sgRNA-target plasmids and donor templates containing the desired mutations into nos-Cas9 embryos. Following multiple rounds of genetic crosses using balancer chromosomes, flies carrying the targeted mutations were identified via PCR and confirmed by sequencing. Assays were performed at two points: mid-age (3 weeks old) and old age (7 weeks old). As previously described^26, 29, 33, 34^, flies were housed in controlled environmental chambers set to 23°C and 50% humidity under a light-dark cycle lasting 12 hours each. Adult flies were collected immediately after eclosion, sorted by sex, and housed in groups of about 10 per vial. They were maintained in cornmeal diet fed ad-libitum and moved to vials with fresh food every 4 to 7 days until the experimental endpoints.

### 2. Skeletal Muscle and Locomotor Performance

To evaluate the functional effect of MADD-associated mutations on muscles, flight ability and negative geotaxis behavior was assessed in adult flies at 3 weeks and 7 weeks.

#### a. Flight Performance

The flight assay was adapted from established protocols to assess age- and genotype-dependent motor performance^26, 35^. Briefly, groups of 15–30 adult *Drosophila* were gently released into the center of a vertically oriented Plexiglass flight chamber illuminated from above. Individual flies were scored based on their directional flight response: upward (score = 6.0), horizontal (4.0), downward (2.0), or flightless (0.0). Flight Index (FI) was calculated for each cohort, representing the average flight performance of the group. Comparisons were made across genotypes and age groups, with all experiments conducted in parallel with age-matched control lines. Detailed information on fly age, genotype, experimental conditions, number of cohorts, total flies tested, and cohort-wise FI values is provided in the Source Data file.

#### b. Geotaxis Performance

As described in previous studies^26, 36^, flies were transferred to a fresh vial containing 8–12 flies per trial. A custom-built 3D-printed device was used to hold up to 12 vials simultaneously. This setup included a Raspberry Pi Camera precisely aligned with the vials, connected to a Raspberry Pi 4B and monitor. After a 2-minute acclimation period, the vial was tapped three times to initiate a negative geotaxis response. Climbing behavior was recorded on video for subsequent analysis using a Faster R-CNN deep learning model to detect vial boundaries. Machine learning algorithms were then used to quantify climbing distance, velocity, and performance indices using methods developed by our group^37^. 3-5 cohorts per genotype and age were included in the assay totaling 50-60 flies.

### 3. Sleep-Circadian Activity Analysis

Sleep-wake behavior and circadian activity in Drosophila were assessed using the Drosophila Activity Monitoring (DAM) system (TriKinetics Inc., Waltham, MA) under controlled 12-hour light:12-hour dark (12L:12D) cycles at 25°C and 50–70% humidity. Individual flies were placed in glass tubes containing standard cornmeal food and allowed to acclimate for two days prior to data collection. Activity was recorded as infrared beam crossings, representing locomotor events. Sleep was defined as any period of at least five consecutive minutes without beam breaks. Flies that did not survive the full monitoring period were excluded from analysis. All metrics were represented as averages over the respective monitoring periods. Sleep metrics including total sleep, daytime sleep (ZT0–ZT12), nighttime sleep (ZT12–ZT24), sleep bout number, bout length, and sleep fragmentation (defined as the number of 1-minute wake episodes per 24 hours) were quantified using a proprietary one-click app developed by our lab using machine learning.

### 4. Cytological Analysis

Like our recently published protocol^33, 38^, fly thoraces were fixed in 4% paraformaldehyde (PFA) in PBS for 15 minutes at room temperature, followed by three 10-minute PBS washes. Samples were embedded in OCT, frozen at −20 °C, and cryosectioned at 20 μm for heart and 30 μm for thorax. Sections were taken at enough depth to reveal the IFMs of thorax. Flies were embedded by orienting them on their dorsal side for the former and lateral side for the latter(sagitally). Each slide included samples from all genotypes and experimental groups within a cohort to minimize inter-cohort and inter-slide staining variability. Slides were washed (3 × 10 min in PBS) to remove residual OCT and then stained with Phalloidin-488 and Lipid-Spot-610 for 30 minutes. It was followed by (3 × 5 min) PBS wash. Slides were mounted in the Vectashield mounting media with DAPI. Immunofluorescence imaging was collected using an Olympus BX-63 microscope at 10× magnification, capturing FITC, TRITC, and DAPI channels. Fluorophores were selected based on their emission spectra and minimal spectral overlap to ensure precise signal discrimination. Imaging was performed on randomly selected, non-overlapping fields within each sample. To ensure representativeness, 3-5 samples per experimental group were imaged and analysis was done with the help of CellSens software by quantifying lipid intensity.

### 5. Measurement of Reactive Oxygen Species (ROS)

ROS levels were assessed with the help of previously published methods^34^ using Dihydroethidium (DHE) staining. 3-week male flies were dissected to isolate thoraces and then submerged in Schneider’s medium to maintain tissue hydration. Thoraces were then embedded in OCT without sucrose or fixation to preserve native redox state and cryosectioned onto slides. Sections were washed once in PBS for 10 minutes, followed by incubation with a 10 µM DHE working solution prepared in Schneider’s medium for 15 minutes at room temperature in the dark. After a 10-minute PBS wash, sections were mounted in Vectashield medium and imaged immediately under identical acquisition settings.

### 6. Cardiac Physiology Analysis

*Ex vivo* physiological analysis of semi-intact *Drosophila* hearts was performed following established protocols^30, 39^. Heart contractions were recorded using direct immersion optics paired with a high-speed digital camera (Hamamatsu Flash 4) capturing at 200 frames per second. Thirty-second videos of beating hearts were acquired using HC Image software (Hamamatsu). Cardiac function was evaluated from high-speed video recordings using the Semi-Automated Optical Heartbeat Analysis (SOHA) software, which quantifies parameters such as heart rate, heart period, systolic and diastolic diameters and intervals, rhythmicity index, and FS^40^. The software also generates mechanical-mode traces for detailed visualization of heart dynamics^40^. 25-35 hearts from male flies at 3 and 7 weeks of age were analyzed for each genotype.

### 7. Measurement of ATP

ATP content in the fly thorax was quantified based on published methods^36, 41^. 8-10 flies from each group were dissected and the thoraces were placed in cold PBS in ice. Tissues were homogenized in RIPA buffer containing protease inhibitor using a pellet pestle while keeping samples chilled throughout. The homogenate was then centrifuged at 16,000 × g for 10 minutes at 4 °C, and the clear supernatant was transferred to a fresh, pre-chilled tube for downstream ATP quantification and protein measurement. Luciferin and luciferase reagents were prepared according to the manufacturer’s instructions. ATP standards were generated from the kit stock to produce a multi-point standard curve on the same plate. Samples and standards were loaded into opaque 96-well plates, the reaction mix was added, and luminescence was read at 560 nm using Synergy LX Multi-Mode Reader (Agilent BioTek Instruments). Each group was assayed in technical triplicate, with three biological replicates per genotype. ATP values were interpolated from the standard curve and normalized to protein concentration for each sample. All readings were collected under identical instrument settings, and background signal was subtracted using reagent-only wells

### 8. Gene Expression Analysis

Gene expression analysis was performed using Real Time-qPCR as previously reported by our group for *Drosophila* tissues^33^. The flies were dissected to isolate thorax and flash frozen in liquid nitrogen and conserved in −80 °C until use. Total RNA was extracted from each biological replicate containing 3-4 thoraces using Zymo Quick-RNA MicroPrep Kit (R1051, Zymo Research, Irvine, CA, USA). Briefly, tissues were homogenized in 150 µL of lysis buffer, incubated at room temperature, and passed through spin columns for purification. Genomic DNA was removed by DNase I treatment (15 min), and RNA was eluted in RNase/DNase-free water. RNA concentration and purity were determined using a Synergy LX Multi-Mode Reader (Agilent BioTek Instruments). Complementary DNA (cDNA) was synthesized from 500 ng of total RNA using iScript RT Supermix (Bio-Rad, 1708840). Reverse transcription was performed under the following conditions: priming at 25 °C for 5 min, reverse transcription at 46 °C for 20 min, and enzyme inactivation at 95 °C for 1 min. qPCR was conducted using 5 ng of cDNA, 200 ng of gene-specific primers, and SsoAdvanced Universal SYBR Green Supermix (Bio-Rad, #1725275) on a CFX Opus Real-Time PCR System (Bio-Rad). Expression levels were normalized to Rpl11 (60S ribosomal protein). Relative gene expression was calculated using the 2^−ΔΔCt method, normalized to reference genes and control samples. All reactions were performed in technical duplicates. Genes with corresponding forward and reverse primers used in this study are: Cpt1-F: TTGCCATCACCCATGAGGG; Cpt1-R: CAATCGCTTTTTCCAGGAACG; SdhA-F: TCGGTGGACAGAGTCTGAAGT; SdhA-R: GCAGCGAGTGACCAGTACG; Dsrebp-F: AGTCGCCGCTTCTCGTCTA; Dsrebp-R: TGTATGGTGGCTGTTGGTTGG; Mef2-F: GAAGCCGAAACGGACTACACA; Mef2-R: GTTGTCGCCGTAAGATCCCG; Duox-F: ATGGCTGGTACAATAACCTGGC; Duox-R: AACCCCATCCGAATAGGAGGG; Grim-F: CAATATTTCCGTGCCGCTGG; Grim-R: CGTAGCAGAAGATCTGGGCC; Ter94-F: AGTCGCGGTGTCCTTTTCTAC; Ter94-R: GGACCCTTGACTGAGATGAAGTT; OPA1-F: CGAGGAGTTCCTACTTGCCG; Opa1-R: GTATCGCAGCTTGAGGGCTC; Pgc-1α (Srl)-F: TATGGAGTGACATAGAGTGTGCT; Pgc-1α (Srl)-R: CCACTTCAATCCACCCAGAAAG; Ampkα-F: TGGGCACTACCTACTGGG; Ampkα-R: ATCTGGTGCTCGCCGATCTT; EtfQO-F: TGCTGTCTTCGGTAGCACTG; EtfQO-R: GTTGATGTTCTGCTTGGGGT; ATPsynD-F: AGTACGAAGCCCTGAAGGTG; ATPsynD-R: CAGAGACTTGAGATGGGCGAT; ATPsynC-F: CCACAGATCAGGTCATTCCAGA; ATPsynC-R: CGAATACTGTTCCGATACCAGC; Tfam-F: AACACCCAGATGCAAAACTTTCA; Tfam-R: GACTTGGAGTTAGCTGCTCTTT; Hid-F: CACCGACCAAGTGCTATACG; Hid-R: GGCGGATACTGGAAGATTTGC; Reaper-F: TGGCATTCTACATACCCGATCA; Reaper-R: CCAGGAATCTCCACTGTGACT; Cox5a-F: CAAAAGGCCACCCTCTATCCC; Cox5a-R: GCATCAATGTCTGGCTTGTTGAA; Etfqo_1 F: TGCTGTCTTCGGTAGCACTG; Etfqo_1 R: GTTGATGTTCTGCTTGGGGT; Etfqo_2-F: TGAAAATGCACAGAGTTCGGAG; Etfqo_2-R: ATAGTGGGTGGTTATCCTGGG.

### 9. Exercise Protocol in Drosophila

As depicted in the schematic in the result section, 3 days old flies were randomly assigned to exercise or sedentary cohorts and subject to negative geotaxis-based endurance training system, which exploits the innate climbing response of flies^42^. We designed and 3D-printed an exercise setup which holds up to 10 *Drosophila* vials simultaneously. The flies were housed in standard vials and placed in the exercise apparatus, which allows an operator to repeatedly drop vials to a flat surface, prompting the flies to climb upward. Each session consisted of 15– 20 drops per minute for 15 minutes daily, five days per week. All exercise sessions were conducted at 1 PM, corresponding to approximately ZT6 under the 12:12 light–dark cycle. The exercise duration and drop frequency was selected based on empirical observation of tolerance and effect of survival on the mutant flies. After each session, flies were returned to fresh food vials. All experiments were performed in parallel across genotypes and sexes.

### 10. Real-time Respiration

Oxygen consumption was measured to assess mitochondrial function using a fluorescence-based optical microplate system (Loligo Systems, #SY25200), which enables real-time, non-invasive monitoring of respiration^43^. Briefly, flies were transferred into 80 µL glass chambers mounted on a 24-well microplate placed atop a 24-channel optical fluorescence oxygen reader, and the entire setup was maintained at 37 °C in a temperature-controlled incubator. Oxygen concentration was recorded at defined intervals for up to 60 minutes, and blank wells containing no flies served as controls for baseline correction. Respiration rates were calculated as the decline in oxygen concentration (mmol/L per unit time) after subtracting blank values.

### 11. Statistical Analysis

Statistical analyses were performed using GraphPad Prism version 10 and Python. Assays were analyzed using either one-way or two-way ANOVA, depending on the experimental design. Post hoc comparisons were conducted using Tukey’s, Dunnet’s or Dunn’s multiple comparisons as appropriate. Data are presented as mean ± standard deviation (SD), and statistical significance was defined as follows: p < 0.05 (*), p < 0.01 (**), p < 0.001 (***), and p < 0.0001 (****). Raw data along with detailed statistical comparisons across genotypes and age groups are provided in the Source Data file.

## Results

### 1. CRISPR Knock-In Etf-QO Mutants Exhibit Impaired Skeletal Muscle Performance and low oxygen consumption

We generated CRISPR knock-in *Drosophila* models carrying three Etf-QO missense mutations L127R, S296C, and L399F within conserved FAD and ubiquinone binding domains, recapitulating late-onset MADD variants. These lines were produced by microinjecting sgRNA plasmids and donor templates into nos-Cas9 embryos **(SI1A)**, followed by balancer-based screening and sequence confirmation **(SI1 B-D)**. To assess skeletal muscle performance, we evaluated skeletal muscle performance using flight ability of the flies at 3 and 7 weeks. All Etf-QO mutants showed reduced flight performance compared with controls, with a stronger deficit at 7 weeks **(Fig. 1C, D)**. Across the mutants, L127R showed the largest decline, S296C displayed an intermediate reduction, and L399F was the least affected, and this order was consistent across sexes and ages. To complement these functional readouts, we measured whole-body oxygen consumption as an index of metabolic capacity. Oxygen consumption measured at 3 weeks was reduced in all three male mutants and in L127R and S296C females **(SI2 A, B)**. The direction of change matched the severity trend observed in flight assays, with L127R showing the largest reduction respiration. As sustained flight in *Drosophila* relies heavily on mitochondrial ATP production, the lower respiration rates provide a metabolic correlate to the reduced flight performance.

**Figure 1.**
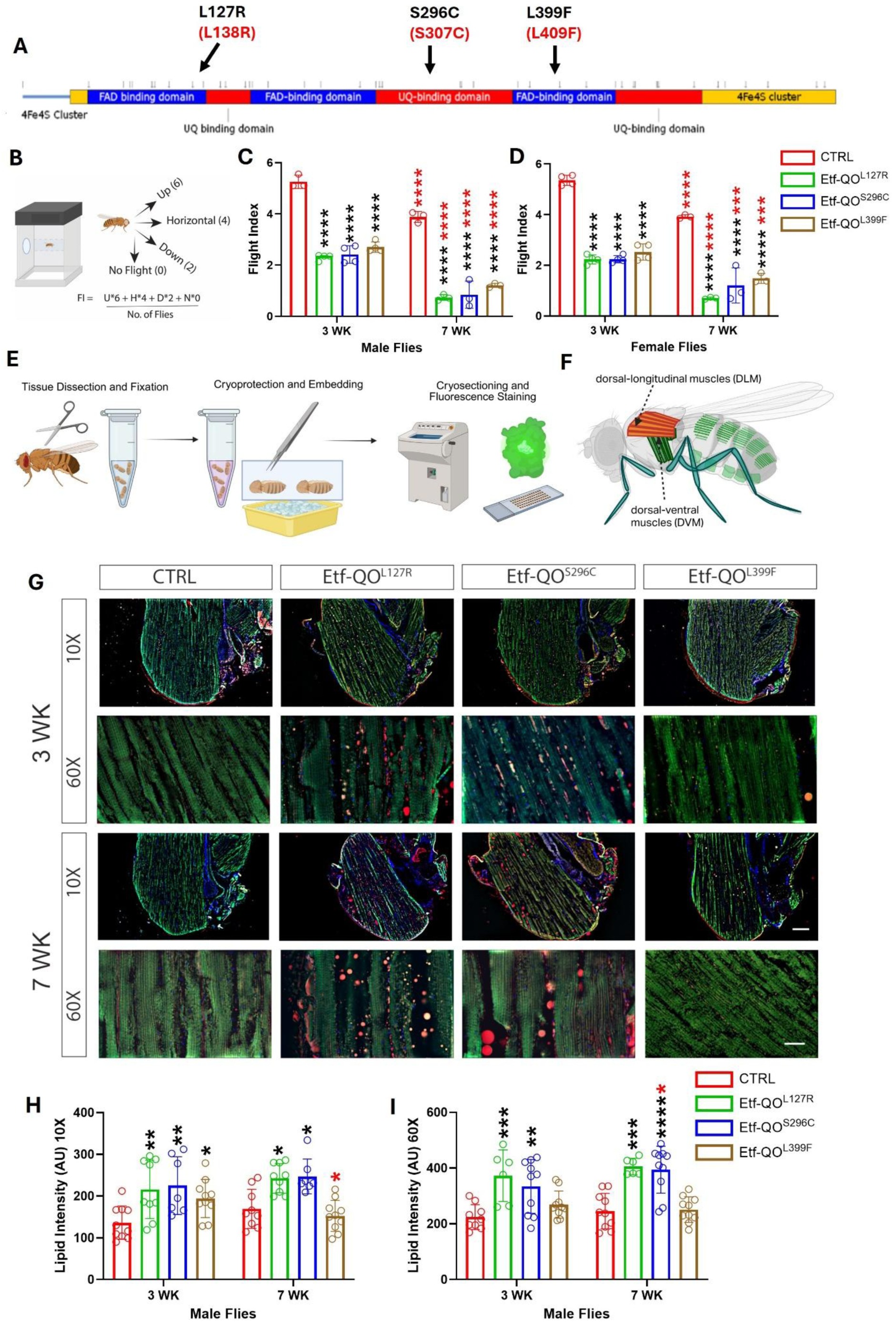
Progressive decline in flight ability accompanied by increased lipid deposition in Etf-QO mutants. Domain architecture of electron transfer flavoprotein-ubiquinone oxidoreductase (Etf-QO) with sites of *Drosophila* (black) and human (red) missense mutation **(A)**. The protein contains two FAD-binding domains (blue), two UQ-binding domains (red), and two 4Fe–4S clusters (yellow) located at the N- and C-terminal regions. Domain arrangement is shown in linear order from N-terminus to C-terminus. Schematic representation of the flight assay setup **(B)**. Flies were released in a vertical flight chamber, and their FI (FI) was calculated and quantified based on their landing positions. The mutants showed diminished flight performance compared to the age-matched controls across both sexes which worsened with age **(C, D)**. n=80–90 flies, 3–5 cohorts. Procedure for cryoimaging IFM muscle **(E)**. Visualization of IFMs in *Drosophila* **(F)**. Thorax and abdomen were isolated by dissecting away wings, legs, and head. Following fixation, tissues were embedded in OCT, cryosectioned to obtain thorax sections, and then subjected to sequential washes and fluorescence staining. Thorax sections were sagittally cryosectioned and stained for visualizing muscle architecture and lipid deposition in IFM **(G).** Lipid droplets (red puncta) were more evident in significantly greater intensity in most of the mutants compared to control in both 3-wk and 7-wk flies **(H, I)**. Phalloidin (green) and DAPI (blue) mark the cytoskeleton and nuclei respectively. Quantification of lipid intensity (n=10–20 images, 5 males/group) confirmed these effects. Two-way ANOVA with Tukey’s multiple comparisons test, *p<0.05, **p<0.01, ***p<0.001, ***p<0.0001. Scale bar: 100 µm (10X), 20 µm(60X). Black and red asterisks denote comparison across genotypes and age respectively. All the raw data associated with this figure are presented in the Source Data file.

### 2. Mutant Flies Exhibit Ectopic Lipid Deposition

Lipid imaging of thoracic tissue revealed increased lipid droplet intensity in all Etf-QO mutants relative to controls at both 3 and 7 weeks (**Fig. 1E–I**). L127R and S296C showed the highest lipid burden across ages, while L399F displayed a milder increase that was most evident at 3 weeks **(Fig. 1H–I, SI3A-C)**. Both male and female flies showed similar genotype-dependent trends, although the intensity appeared slightly elevated in females especially in 7-week L399F **(SI3A-C)**. Because statistical comparisons between males and females were not performed, these observations are presented descriptively. Elevated lipid signal was also confirmed at higher magnification (60×) **(Fig. 1I)**. Thoracic sections were obtained at comparable depths to consistently expose the IFMs. Overall, we observed that ectopic lipid deposition is present across Etf-QO mutation backgrounds, with the strongest signal in L127R and S296C and a comparatively mild effect in L399F.

### 3. Etf-QO Mutations Impair Locomotor Performance and Alter Activity-Sleep Behavior

Unlike flight, which is driven by the IFMs of the thorax, climbing in *Drosophila* is primarily done through leg musculature and neurons. These muscle groups are also distinct in that flight muscles generate high-power, oscillatory output whereas leg muscles perform occasional whole-body locomotion. All Etf-QO mutants showed reduced climbing performance measured with geotaxis assay **(Fig. 2A)**, compared with controls **(Fig. 2C–H)**. L127R consistently exhibited the strongest impairment in both sexes. S296C showed reduced climbing in males at 3 weeks and in females at 7 weeks, while L399F displayed milder deficits. More specifically, the distance climbed at t = 6 seconds was used as a summary metric **(Fig. 2 G, H)** for comparing climbing activity across and showed similar genotype-dependent trend observed in full geotaxis traces. Several deficits in control as well as L127R and L399F were significantly higher in older females indicating age- and sex-dependent variation in locomotor output. Together, these results show that Etf-QO mutations impair leg-driven locomotor behavior in addition to the flight-related deficits reported above. Continuous activity monitoring using the Drosophila Activity Monitoring (DAM) system **(Fig. 2B)** revealed that Etf-QO mutations show altered spontaneous activity and sleep behavior **(Fig. 2I-Q)**. Daytime activity and total activity were significantly reduced in L127R and S296C mutants, whereas L399F exhibited comparatively milder alterations **(Fig. 2I, K)**. Reduced activity was accompanied by increased daytime, nighttime, and total sleep **(Fig. 2L-N)**, consistent with reduced behavioral output. Sleep architecture analysis further revealed genotype-dependent alterations in bout number and bout length **(SI6A-F)**, while sleep-fragmentation measures were elevated in mutant backgrounds **(Fig. 2O-Q)**. Together, these findings suggest that Etf-QO dysfunction affects not only locomotor activity but also sleep organization and consolidation.

**Figure 2.**
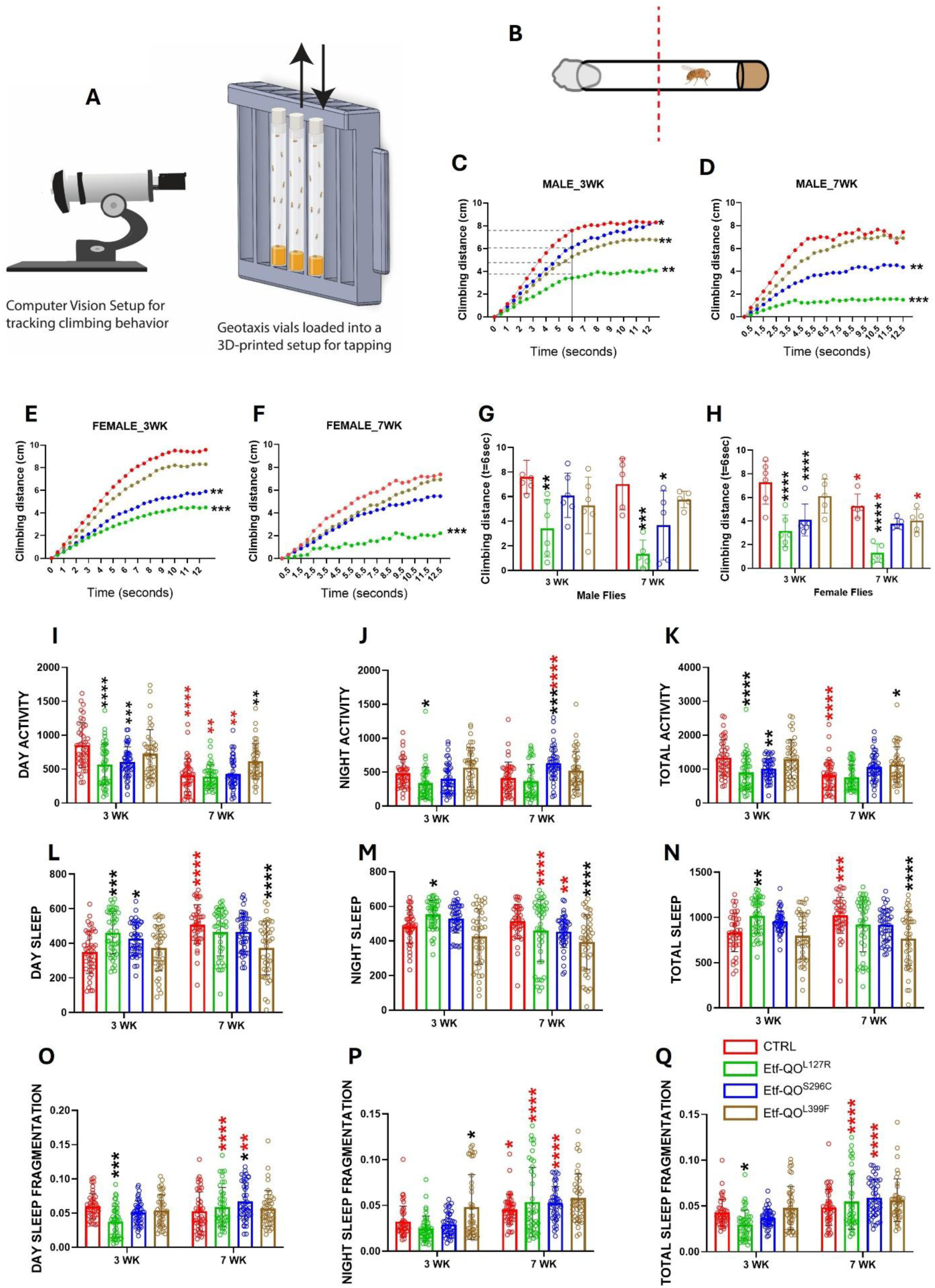
Etf-QO mutations impair locomotion and alter sleep architecture. Schematic of automated geotaxis assay with Raspberry Pi Camera**(A)**. Schematic representation of the Drosophila Activity Monitoring (DAM) system with infrared-beam-based detection of spontaneous locomotor activity and sleep behavior in individual flies (B). Geotaxis traces from male 3-week (C), male 7-week (D), female 3-week (E), and female 7-week (F) cohorts demonstrate reduced climbing performance in mutants (Etf-QO ^L127R^, Etf-QO ^L399F^) across both sexes. genotype-dependent reductions in climbing performance. Quantification of climbing distance at 6 seconds for males (G) and females (H) confirms impaired locomotor function. DAM analysis revealed reduced daytime activity (I), altered nighttime activity (J), reduced total activity (K), increased daytime sleep (L), increased nighttime sleep (M), and increased total sleep duration (N), particularly in L127R and S296C mutants, indicating altered activity-rest behavior. One-way ANOVA with Dunn’s multiple comparisons test, *p<0.05, **p<0.01, ***p<0.001. Distance covered at half duration (t=6 sec) showed significant deficiency Etf-QO L127R females in mid-age as well as old-age **(G)**. Similar trend was observed in male mutants **(F)**. n=50-60 flies, 3–5 cohorts. Two-way ANOVA with Tukey’s multiple comparisons test, *p<0.05, **p<0.01, ***p<0.001, ***p<0.0001, black and red asterisks denote comparison across genotypes and age respectively. All the raw data associated with this figure are presented in the Source Data file.

### 4. Cardiac Performance Deficits in Etf-QO mutants

**Fig. 3 A** outlines the semi-intact heart preparation and cardiac parameters extraction from high-speed videos recordings. Only male hearts were quantified for all cardiac measurements. At 3 weeks, Etf-QO mutants showed cardiac abnormalities relative to controls, including higher Arrythmia Index (AI) across all three mutant lines in both age groups **(Fig. 3 D)**. AI increased significantly in all lines at 7 weeks, with a greater rise in the Etf-QO mutants compared with controls. Fractional Shortening (FS) was also significantly reduced in all the mutant lines at 3 weeks and in L127R and S296C at 7 weeks with 7-week flies exhibiting significantly higher impairment compared to 3-week flies across all lines **(Fig. 3 I)**. Diastolic diameter remained comparable between 3-week and 7-week cohorts, whereas systolic diameter decreased in L127R, S296C, and control **(Fig. 3G-H).** This pattern indicates that the age-related decline in FS and contractile performance is driven primarily by reduced systolic displacement rather than diastolic dilation. Conduction timing parameters showed a comparable pattern among genotypes at 3 weeks, and heart rate declined in all lines by 7 weeks **(Fig. 3B)**. In addition to these cardiac changes, abdominal sections surrounding the heart showed greater lipid accumulation in the fat bodies of 3-week male mutants, with quantitatively elevated levels in L127R and S296C **(Fig. 3 J-K)**, indicating that systemic lipid dysregulation parallels the observed cardiac defects.

**Figure 3.**
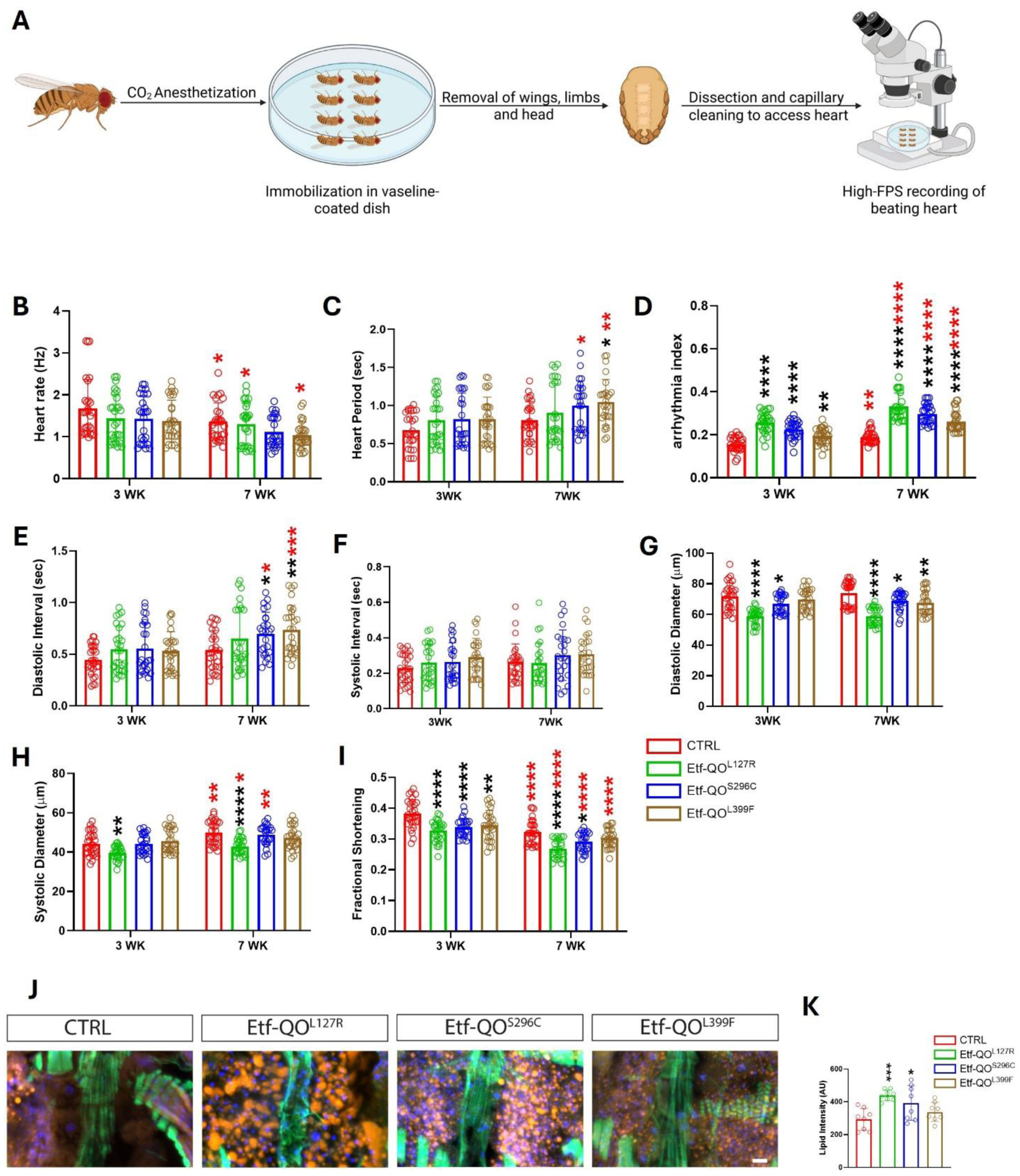
MADD-induced LSM disrupts cardiac performance and lipid accumulation in the fat bodies. Schematic of procedure for visualizing cardiac physiology in *Drosophila* **(A)**. Decreased heart rate and elongated diastolic interval observed in 3-wk male mutants compared to age-matched controls **(B, E)**, which worsen with age. AI **(D)** was higher in Etf-QO ^L399F^ at 3 wk, with significant worsening observed at 7 wk in all three mutants. Two-way ANOVA with Tukey’s multiple comparisons test. *p<0.05, **p<0.01, ***p<0.0001. Black and red asterisks denote comparison across genotypes and age, respectively. n=25-30 flies per genotype and age. Higher lipid accumulation was observed in 3-week male mutants in the fat bodies region surrounding the heart **(J)** and exhibited significantly higher intensity in two mutants **(K)**. One-way ANOVA with Dunnett’s multiple comparisons test, *p<0.05, ***p<0.001, ****p<0.0001, n=8 images, 5 3-week males per genotype. Scale bar: 20 µm. All the raw data associated with this figure are presented in the Source Data file.

### 5. Etf-QO Mutations Increase ROS with Reduced ATP and Altered Mitochondrial and Stress Transcripts

DHE staining of IFM tissue revealed significantly elevated ROS levels in all Etf-QO mutant backgrounds compared with controls **(Fig. 4A-B)**. Consistent with this, expression of the ROS-generating enzyme Duox was elevated across all mutant lines **(Fig. 4L)**. Interestingly, elevated ROS was accompanied by elevation of the mitochondrial antioxidant enzyme SOD2 across all mutant backgrounds **(Fig. 4M)**. However, Nrf2 transcript abundance remained largely unchanged across genotypes **(Fig. 4N)**, despite elevated ROS levels, suggesting that the antioxidant adaptation may occur through post-transcriptional regulation of redox signaling or through Nrf2-independent stress-response mechanisms. Cellular ATP content was significantly reduced in L127R and S296C and relatively preserved in L399F **(Fig. 4C)**. In response to the energetic stress, compensatory mitochondrial remodeling pathways appeared activated. Ampk and Srl (PGC-1α) were significantly upregulated in all mutant backgrounds **(Fig. 4E-F)**. Downstream mediators of mitochondrial remodeling, including Tfam and Opa1, were elevated in selected alleles **(Fig. 4H, G)**, suggesting engagement of mitochondrial biogenesis, mtDNA maintenance, and mitochondrial network remodeling. Pink1 and Parkin transcripts were significantly elevated in L127R and S296C mutants **(Fig. 4J-K)**, suggesting mitochondrial turnover. Additional metabolic transcripts further highlighted allele-specific consequences of Etf-QO dysfunction. Although AtpsynD remained unchanged **(Fig. 4D)**, AtpsynC was significantly reduced in L127R **(SI4A)**. Similarly, Mef2, Cpt1, and SdhA were reduced in L127R (**SI4B-D)**, suggesting impaired muscle maintenance, fatty-acid utilization, and mitochondrial metabolism. Ter94 showed only modest changes **(SI4E)**, indicating limited activation of broader proteostasis pathways.

**Figure 4.**
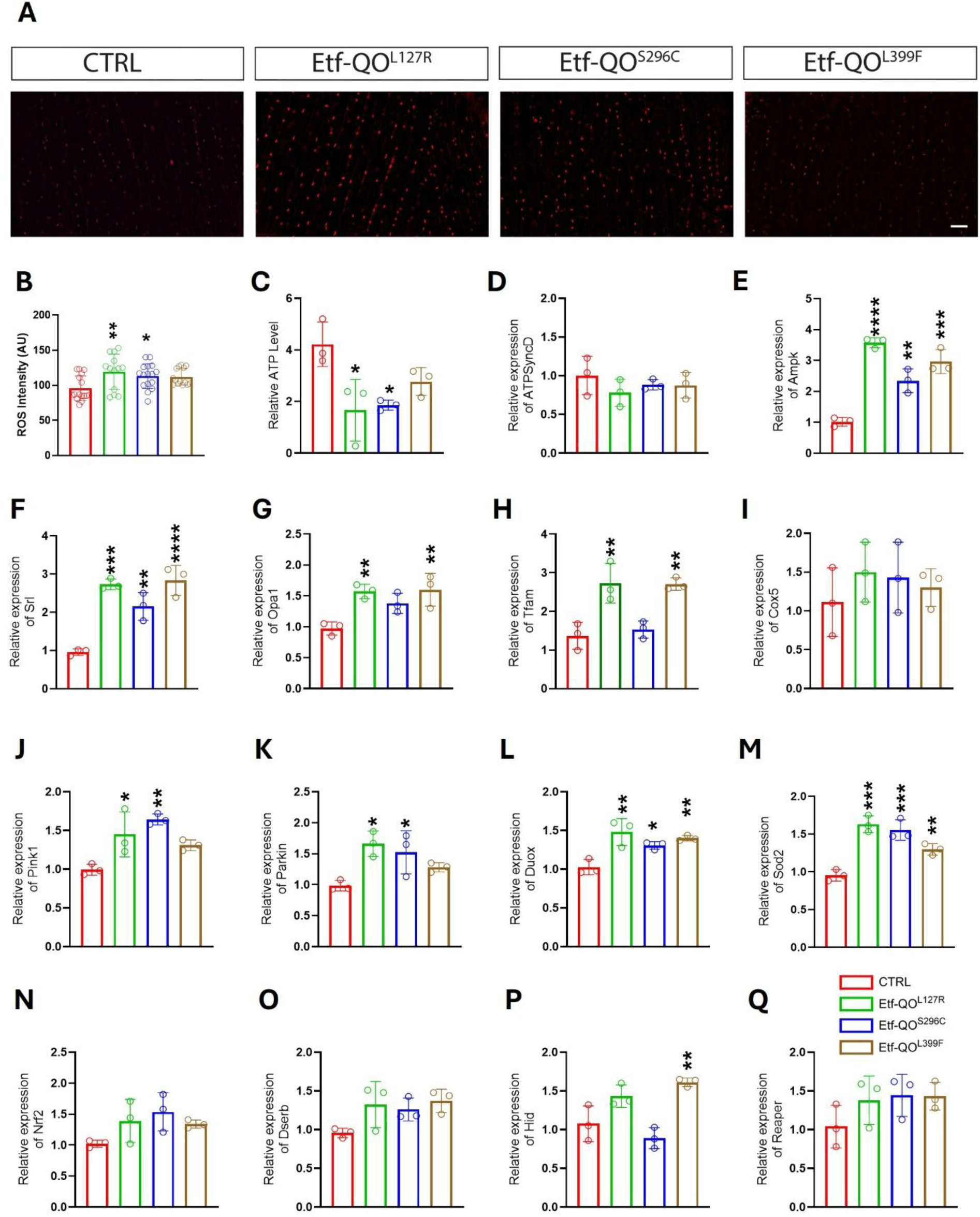
Etf-QO Mutations Elevate ROS Levels and Modify Mitochondrial and Stress-Related Gene Expression. Quantification of ROS using DHE staining **(A, B)**. ATP content was measured from thoraces of adult *Drosophila* melanogaster to assess mitochondrial bioenergetic function at the tissue level **(C)**. Relative mRNA expression levels of genes involved in mitochondrial bioenergetics and remodeling (D-I), mitochondrial quality control (J-K), oxidative-stress responses (L-N), and downstream lipid metabolism and apoptosis pathways (O-Q). Panels J-K represent the mitochondrial quality-control genes Pink1 and Parkin, panels L-N represent oxidative-stress-related genes including Duox, Sod2, and Nrf2, and panels O-Q represent downstream lipid-metabolism and apoptosis-associated transcripts. Expression normalized to Rpl11; data represent mean ± SD from three biological and two technical replicates obtained from thorax samples of 3-week-old males. One-way ANOVA with Dunnett’s multiple comparisons test, *p<0.05, **p<0.01, ***p<0.001, ***p<0.0001. All the raw data associated with this figure are presented in the Source Data file.

### 6. Exercise Mitigates Physiological and Metabolic Deficits and Induces Compensatory Transcriptional Response

Only 3-week males were used to assess the effect of exercise. Exercised cohorts showed improved FI than their sedentary counterparts in all three Etf-QO mutant lines with L127R displaying significantly increased FI **(Fig. 5B)**. In contrast, geotaxis outcomes showed minimal improvement with exercise **(Fig. 6)**. L127R displayed a slight upward shift in climbing performance **(Fig. 6E-F)** in both males and females but this change was not statistically significant, and S296C and L399F showed no clear improvement **(Fig. 6E-F).** These findings indicate that the exercise regimen has minimal effect on leg-driven locomotor behavior although it produced measurable benefits in high-power flight performance. In parallel with the improvement in flight ability, the 15-minute daily training protocol reduced lipid accumulation in dorsal-longitudinal muscles across all mutants **(Fig. 5C–D)**. The reduction was also evident at higher magnification **(SI5)**. DHE imaging revealed a downward trend in ROS across exercised groups, with S296C showing a significant reduction **(Fig. 5E–F)**, suggesting partial alleviation of oxidative stress. Transcriptional analysis showed coordinated activation of mitochondrial remodeling pathways. Tfam and AMPKα increased in all exercised groups, and PGC-1α was significantly elevated in L127R **(Fig. 5G–I)**, supporting engagement of an AMPK-PGC-1α–dependent compensatory program. Together, these findings indicate that moderate endurance exercise reduces lipid and ROS burden, improves muscle performance, and activates mitochondrial biogenesis pathways specifically under Etf-QO dysfunction, consistent with functional and metabolic rescue rather than enhancement of baseline physiology.

**Figure 5.**
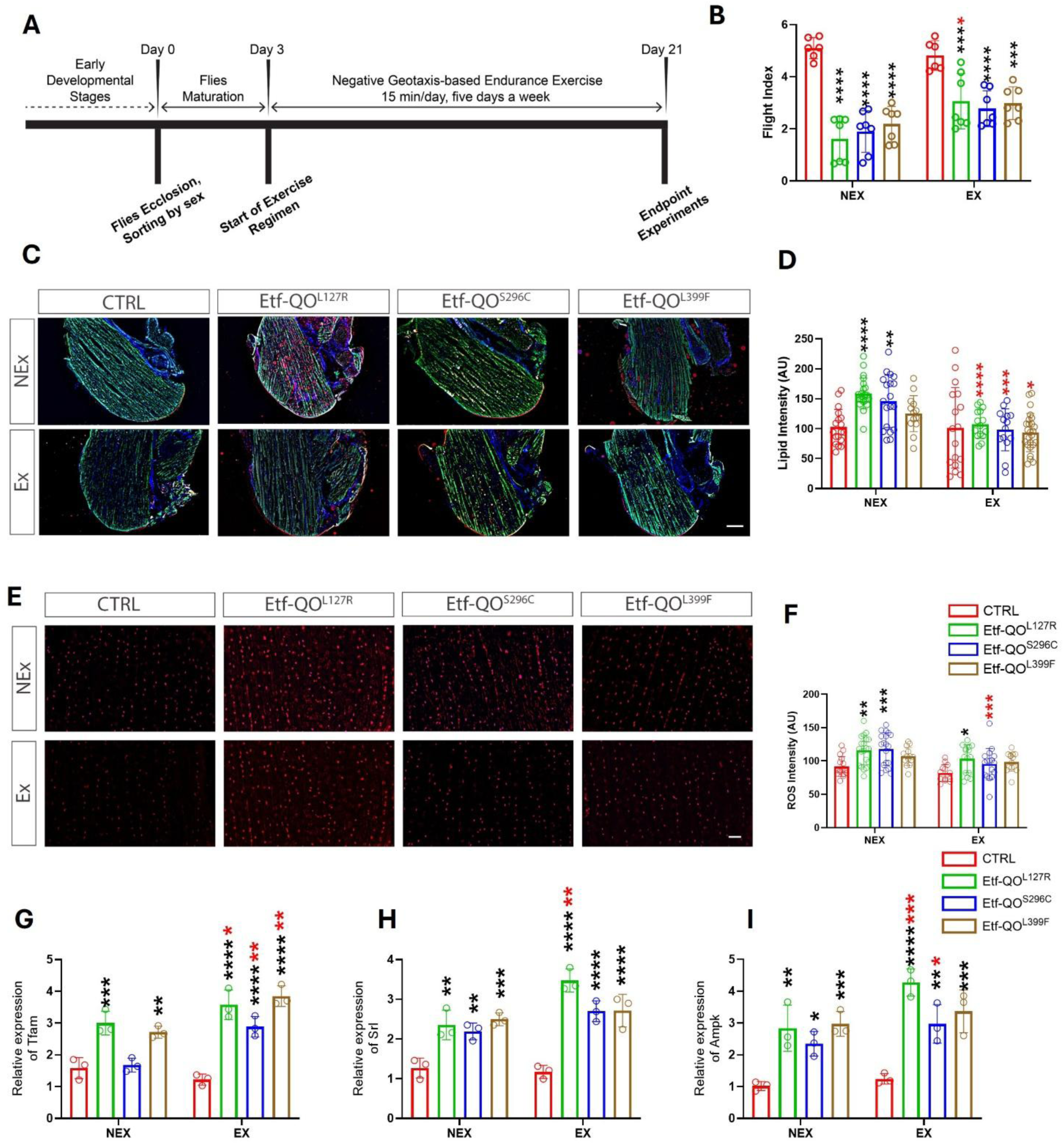
Moderate daily exercise intervention showed beneficial effect in mitigating lipid deposition and improving flight ability. Schematic for implementation of exercise regimen **(A)**. Exercise (15 min/day for 2.5 weeks) improved flight performance compared in 3-week males compared to the sedentary controls. (B). n=80–90 flies, 3–5 cohorts. Higher lipid deposition observed in IFM of Etf-QO mutants was also attenuated as evidenced by visible decrease in lipid intensity **(C, D)** in IF images quantified using CellSense Imaging Software, Olympus Life Science. Lipid reduction was accompanied by decrease in oxidative stress as evidence by DHE staining **(E, F)** along with upregulation of key energy-stress-responsive genes Tfam, Srl and Ampk **(G-I)** n=15-25 images from 5 male flies per group at 3 weeks of age. Two-way ANOVA with Tukey’s multiple comparisons test **p<0.01, and ***p<0.001, Black and red asterisks denote comparison across genotypes and exercise conditions, respectively. Scale bar: 100 µm. All the raw data associated with this figure are presented in the Source Data file.

**Figure 6.**
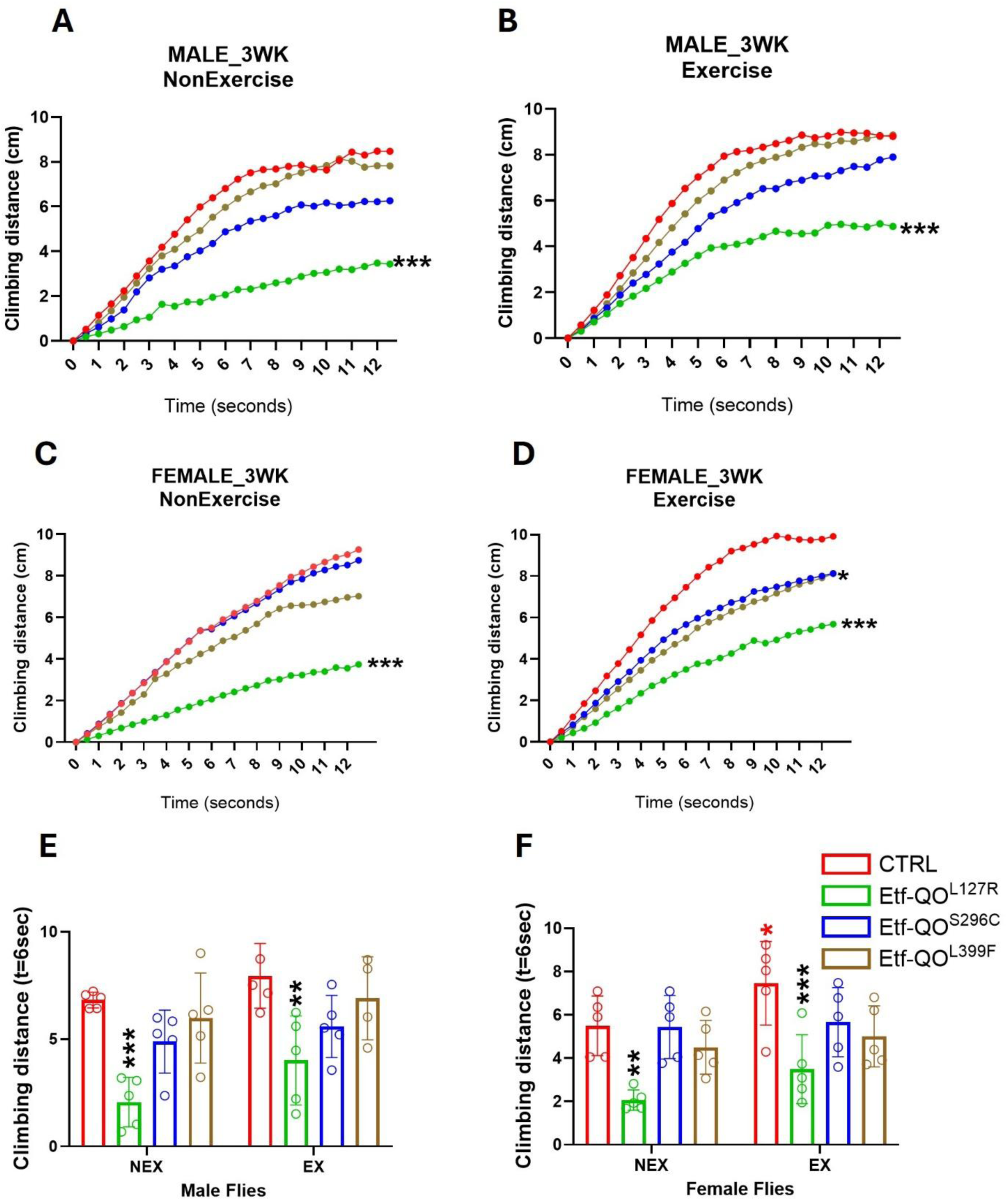
Effect of moderate exercise on locomotor dysfunction linked with Etf-QO mutations. Exercise exhibited mixed results with modest improvement in climbing performance **(A-D)** and distance climbed at half duration in Etf-QO ^L399F^ mutant flies **(E-F)**. Data represent mean ± SD from 3–5 cohorts (n = 80–90 flies per genotype). One-way ANOVA with Dunn’s multiple comparisons test, ***p<0.001 **(A-D)**. Two-way ANOVA with Tukey’s multiple comparisons test, **p<0.01, ***p<0.001. All the raw data associated with this figure are presented in the Source Data file.

### 7. Exercise Improves Cardiac Parameters in Etf-QO mutants

Exercised cohorts showed improved rhythm and contractility within each genotype. AI **(Fig. 7C)** was lower and FS **(Fig. 7H)** was significantly higher in all four lines compared with their sedentary cohorts. Systolic and diastolic diameters **(Fig. E, F)** in exercised groups was associated with a tendency toward smaller end-systolic diameter with little to modest change in end-diastolic diameter, which is consistent with the observed increase in FS. Heart rate was lower and heart period was longer in exercised groups, reflecting improved contraction efficiency **(Fig. 7 A-B)**. Together, these panel-level readouts indicate an exercise-associated shift toward more regular rhythm and stronger contraction across the three Etf-QO mutant lines **(Fig. 7A–H)**.

**Figure 7.**
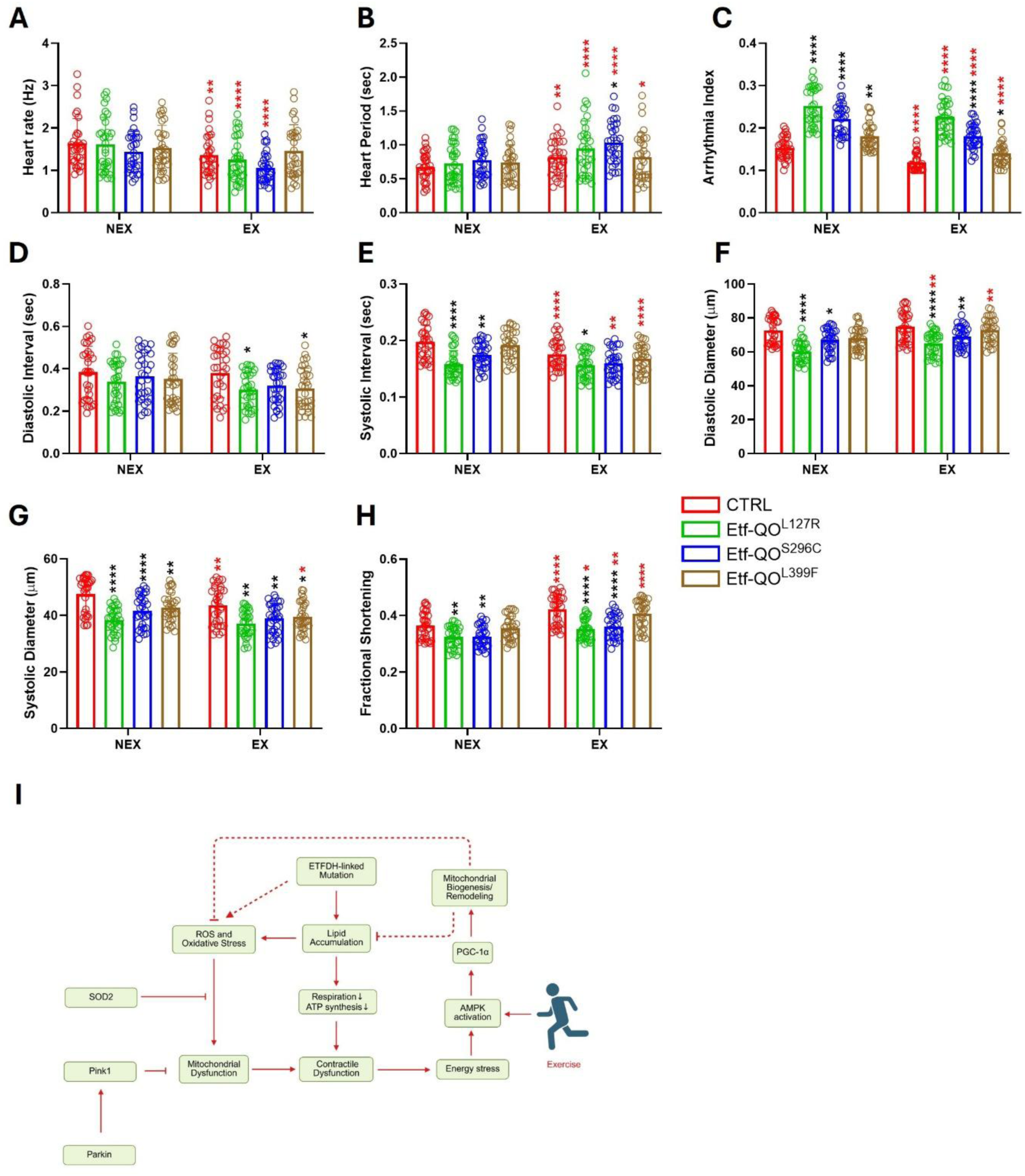
Exercise-induced improvements in cardiac function in Etf-QO mutants and a working model illustrating the underlying metabolic and mitochondrial responses. Sedentary mutants exhibited cardiac dysfunction characterized by elevated AI and reduced FS which was significantly improved in exercised cohorts as evidenced by decreased AI (**C)** and increased FS **(H)** indicating enhanced pumping efficiency. Working model summarizing the relationship between ETFDH-linked dysfunction, mitochondrial stress and exercise-associated adaptive responses **(I)**. Etf-QO mutation promotes lipid accumulation and oxidative stress, which are associated with reduced respiration and ATP synthesis and mitochondrial dysfunction. Mutant flies exhibit activation of energy-stress signaling, including increased AMPK and PGC-1α expression, together with markers of mitochondrial remodeling such as Tfam and Opa1. Elevated SOD2 expression suggests activation of antioxidant defense pathways, whereas increased Pink1 and Parkin expression is consistent with engagement of mitochondrial quality-control mechanisms in response to mitochondrial stress. Exercise is proposed to enhance AMPK–PGC-1α signaling and mitochondrial remodeling, thereby partially counteracting mutation-associated oxidative stress and contractile dysfunction. Solid arrows indicate experimentally observed associations, whereas dashed arrows denote proposed or inferred relationships. n = 25–30 hearts per genotype from 3-week-old male flies. Two-way ANOVA with Tukey’s multiple comparisons test, *p<0.05, **p<0.01, **p<0.001, ****p<0.0001, black and red asterisks denote comparison across genotypes and exercise conditions, respectively. All the raw data associated with this figure are presented in the Source Data file.

## Discussion

### 1. Modeling Human ETFDH Mutations in Drosophila

Clinically, MADD presents with a spectrum of symptoms, including progressive muscle weakness, lipid accumulation in the skeletal and cardiac muscle, and in some cases, cardiomyopathy and metabolic crises^9, 44, 45^. To model MADD phenotypes *in vivo*, we generated CRISPR knock-in *Drosophila* lines carrying patient-relevant missense mutations, L127R, S296C, and L399F, in the Etf-QO gene. Our CRISPR knock-in mutations reflect well-documented pathogenic contexts of human ETFDH variants (**Fig. 1A**). The L127R fly variant maps to the conserved FAD-binding domain position p.Leu138Arg (c.413T>G), reported as pathogenic in ClinVar^46^ and consistently associated with late-onset MADD featuring progressive proximal weakness, exercise intolerance, myalgia, elevated CK, and episodic rhabdomyolysis^15, 47^. The S296C fly variant lies within the FAD domain and parallels human p.Ser307Cys (c.920C>G), classified likely pathogenic^48^ and situated in the riboflavin-responsive mutational spectrum^49–51^. The L399F fly variant corresponds to the UQ-binding domain and aligns with human p.Leu409Phe (c.1227A>C), designated pathogenic^52^ and linked to variable severity with occasional cardiac involvement, often improved by riboflavin and CoQ10^53–55^. Consistent with these human patterns, the three fly lines exhibited progressive locomotors **(Fig. 1C-D)** and cardiac dysfunction **(Fig. 3 D, I)**, lipid accumulation **(Fig. 1 G-I, SI3 A-C)**, and oxidative stress **(Fig. 4 A-B)**. Moreover, L127R mutant showed significant reduced climbing ability across the cohorts **(Fig. 2 C-H).** Other mutant S296C showed impaired locomotor ability at 7 weeks for male **(Fig. 2 D)** and 3 weeks for male **(Fig. 2 E)** suggesting differential effects of the mutations due to allele-specific disruption of Etf-QO enzymatic efficiency and variable thresholds of metabolic stress tolerance across muscle groups. L127R lies within the conserved FAD-binding domain of Etf-QO adjacent to a 4Fe–4S cluster that supports electron transfer from upstream acyl-CoA dehydrogenases. Missense mutation at this position seems to more strongly disrupt cofactor interaction and electron-transfer efficiency. The precise structural basis behind this requires further study. Immunofluorescence imaging revealed ectopic lipid accumulation in thoracic **(Fig. 1 G)** and abdominal tissues **(Fig. 3 J)**, mimicking the lipid accumulation seen in human tissues^56^. Cardiac assessments further demonstrated functional impairments such increased AI **(Fig. 3 D)**, and reduced FS **(Fig. 3 I**), potentially due to metabolic or mitochondrial dysregulation. In fact, consistent with our findings impaired contractile performance is associated with mitochondrial cardiomyopathy reported in MADD^44^. Overall, the L127R mutation produced the most severe phenotypes across muscular, metabolic, and cardiac domains, suggesting that specific mutations may differentially impact Etf-QO function and disease severity^57, 58^. This mirrors the clinical heterogeneity observed in MADD patients, where genotype-phenotype correlations are increasingly recognized^45^. While not a complete representation of the human condition, the model offers a platform for exploring the disease in a controlled setting.

### 2. Progressive Neuromuscular and Behavioral Decline Reflects Clinical MADD Trajectory

In our *Drosophila* models, we observed a clear decline in motor performance that recapitulated the clinical trajectory of late-onset MADD. Flight ability was markedly reduced across all alleles and worsened with age, whereas geotaxis defects were present but less pronounced **(Fig. 1C– D, Fig. 2C–H)**. FI worsened with age, mirroring the progressive motor decline seen in human patients^57^. Sustained flight in flies relies heavily on mitochondrial ATP production, and the combined reduction in FI and oxygen consumption **(SI2)**, together with reduced daytime activity **(Fig. 2I)** and lower total activity **(Fig. 2K)** is consistent with a progressive energetic constraint affecting multiple physiological systems. These patterns indicate that tissues with high energetic demand are disproportionately affected when electron-transfer capacity is compromised and underscore the critical role of Etf-QO gene in maintaining mitochondrial energy homeostasis in high-demand tissues such as skeletal muscle^1^. The gradual decline in motor output is consistent with accumulating metabolic stress in muscle, including impaired fatty-acid oxidation, reduced ATP availability, and accompanying redox imbalance observed in the mutant backgrounds. The Drosophila Activity Monitoring (DAM) system demonstrated that Etf-QO dysfunction influences sleep behavior. Mutant flies, particularly the severely affected L127R line, exhibited increased daytime, nighttime, and total sleep duration together with reduced daytime and total activity **(Fig. 2I-N)**. Because sleep is increasingly recognized as a physiological state that promotes energy conservation, metabolic recovery, and mitochondrial homeostasis^59, 60^, the increase in sleep observed in L127R mutants may represent a compensatory response to chronic bioenergetic stress imposed by Etf-QO dysfunction rather than merely a secondary behavioral deficit. Sleep architecture analysis further revealed genotype-dependent alterations in bout number and bout length **(SI6A-F)**, whereas measures of sleep fragmentation were elevated in mutant backgrounds **(Fig. 2O-Q)**, consistent with reduced sleep consolidation. Notably, the severity hierarchy of these behavioral abnormalities closely mirrored the pattern observed for ATP depletion, respiratory dysfunction, and locomotor impairment, supporting a shared underlying mechanism of mitochondrial dysfunction and chronic energetic stress.

### 3. Ectopic Lipid Deposition in Etf-QO Mutations

A hallmark of MADD is the pathological accumulation of neutral lipids in muscle tissues^1^. In our Etf-QO mutations model, we observed a pronounced level of lipid droplet in the thoracic tissues in the mutants compared to the age- and sex-matched control **(Fig. 1 E-G).** These findings reflect clinical observations in MADD patients, where muscle biopsies often reveal vacuolated fibers filled with lipid droplets, especially in late-onset cases^61^. This phenotype worsened with age although it was statistically insignificant **(Fig. 1 H-I)**. Mechanistically, lipid accumulation likely results from a bottleneck in mitochondrial β-oxidation caused by ETFDH dysfunction, which impairs electron transfer to the respiratory chain and leads to the buildup of acyl-CoA intermediates^57^. These intermediates are then diverted into cytoplasmic lipid droplets **(Fig. 1 G-I)**, a compensatory but ultimately pathological response^62^. In human LSM cases, lipid overload has been implicated in sarcomeric disorganization and muscle fiber degeneration^63^. This deterioration may arise from altered lipid–protein interactions at the sarcomere or from ROS-mediated damage to cytoskeletal components, both of which are exacerbated by mitochondrial dysfunction^63^. Our mutants show both lipid accumulation and elevated ROS **(Fig. 4A–B)**, suggesting that defective fatty-acid utilization and redox stress act together to exacerbate muscle vulnerability.

### 4. Cardiac Dysfunction Linked with Mitochondrial Impairment

Mitochondrial dysfunction is evident in Etf-QO mutants from significantly reduced thoracic ATP in L127R and S296C **(Fig. 4C)** together with a compensatory transcriptional program indicative of energy stress and remodeling, including increased AMPKα and PGC-1α(Srl) **(Fig. 4E-F)** and allele-specific elevations in Opa1 and Tfam **(Fig. 4G-H)**. These mitochondrial abnormalities align with the cardiac phenotype as mutants exhibited hallmark features of cardiomyopathy such as elevated AI and reduced FS **(Fig. 3 D, I)**, together with age-related changes in end diastolic interval **(Fig. 3E)**. Collectively, these data indicate that compromised mitochondrial energetics contribute to cardiac dysfunction in this model.

Clinically, cardiac involvement is a recognized component of MADD, encompassing cardiomyopathy, arrhythmias, and reduced cardiac output ^64^. Our findings in *Drosophila* are congruent with reports that defects within electron-transfer components precipitate bradycardia, rhythm instability, and diminished contractile efficiency in mitochondrial disorders ^65, 66^. Mechanistically, impaired electron transfer between ETF and the respiratory chain can reduce ATP production and increase oxidative stress, processes expected to exacerbate Ca²⁺ handling deficits and sarcomeric dysfunction^65, 67^. Importantly, this work provides one of the first *in vivo* demonstrations of cardiac dysfunction in a genetically engineered *Drosophila* model of patient-relevant ETFDH mutations^68^.

### 5. Redox Imbalance and Compensatory Pathways in Disease Pathogenesis

Oxidative stress can play a major role in pathogenesis and progression of metabolic disorders, including MADD, diabetes, obesity, and metabolic syndrome^69^. All three mutant lines exhibited significantly elevated ROS levels, as evidenced by increased DHE fluorescence intensity **(Fig. 4A-B)**, accompanied by concordant upregulation of the ROS-generating enzyme Duox **(Fig. 4L)**. This oxidative stress likely arises from impaired electron transfer within the mitochondrial respiratory chain due to ETFDH dysfunction, which leads to electron leakage and the generation of superoxide radicals^70^. The accumulation of ROS can damage mitochondrial DNA, proteins, and lipids, further exacerbating mitochondrial dysfunction and contributing to tissue degeneration^70^. These findings are consistent with previous studies in patient-derived cells, where ETFDH mutations have been associated with increased oxidative stress and mitochondrial damage^70^. Our data extends these observations by providing *in vivo* evidence of redox imbalance in a genetically tractable *Drosophila* model, enabling precise dissection of the temporal and tissue-specific dynamics of oxidative stress in LSM.

At the molecular level, the redox imbalance observed in Etf-QO mutants likely acts as both a consequence and a driver of disease progression. Impaired electron transfer through Etf-QO is expected to reduce respiratory efficiency and promote electron leakage, resulting in elevated ROS and diminished ATP production. Chronic oxidative stress can further compromise mitochondrial function by damaging proteins, lipids, and nucleic acids, thereby exacerbating energetic insufficiency and accelerating tissue degeneration^70, 71^. Consistent with this model, transcript analysis revealed activation of a coordinated compensatory response encompassing mitochondrial remodeling, antioxidant defense, and mitochondrial quality control. Spargel (Srl/PGC-1α) and Ampk were upregulated in all three mutant lines **(Fig. 4E-F)**, supporting activation of energy-stress signaling pathways that promote mitochondrial adaptation. Downstream mediators of mitochondrial remodeling, including Tfam and Opa1, were elevated in selected alleles **(Fig. 4 G, H)**, suggesting engagement of mtDNA maintenance and mitochondrial network remodeling mechanisms. These responses are known to preserve mitochondrial integrity and support contractile function in metabolically demanding tissues such as skeletal and cardiac muscle^72–75^. Pink1 and Parkin transcripts were significantly increased in L127R and S296C mutants **(Fig. 4J-K)**. Because the Pink1-Parkin pathway plays a central role in the recognition and removal of dysfunctional mitochondria, these findings suggest activation of mitochondrial quality-control mechanisms in mutant backgrounds experiencing the greatest metabolic stress. Elevated ROS was further accompanied by increased expression of the mitochondrial antioxidant enzyme Sod2 across all three alleles **(Fig. 4M)**, indicating engagement of antioxidant defenses in response to persistent oxidative burden. Interestingly, Nrf2 transcript levels were not significantly altered **(Fig. 4N)**, suggesting that antioxidant adaptation may occur through post-transcriptional activation of redox signaling or through alternative stress-responsive pathways.

Additional transcriptomic changes further reflected disease burden. Cox5 exhibited an upward trend across all mutant backgrounds **(Fig. 4I)**, which may represent an attempt to sustain electron flux and preserve oxidative phosphorylation despite upstream limitations imposed by ETFDH dysfunction^76, 77^. Mef2 expression was reduced in L127R and S296C **(SI4B)**, consistent with impaired myogenic maintenance, whereas the apoptotic regulators Hid and Reaper showed slight activation **(Fig. 4P-Q)** although not statistically significant. Dsrebp also trended upward in all mutants **(Fig. 4O)**, matching the ectopic lipid accumulation observed histologically **(Fig. 1G)**. Notably, the severity pattern observed for Pink1, Parkin, and SOD2 expression closely paralleled the physiological phenotypes, with L127R generally exhibiting the strongest response, S296C showing intermediate effects, and L399F displaying milder alterations. The close correspondence between physiological severity and activation of Pink1, Parkin, and SOD2 suggests that mitochondrial quality-control and antioxidant defenses are recruited in proportion to the degree of metabolic stress imposed by individual Etf-QO mutations. The behavioral abnormalities detected by DAM monitoring further support this interpretation. The same mutant alleles that exhibited the greatest ATP depletion, oxidative stress, and activation of Pink1–Parkin quality-control pathways also displayed the largest reductions in spontaneous activity and the strongest sleep abnormalities **(Fig. 2I-Q; SI6A-F)**.

### 6. Exercise as a Non-Therapeutic Intervention of LSM

A major translational insight from this study is the therapeutic potential of low-to-moderate exercise in mitigating the pathological features of LSM caused by ETFDH mutations. In our models of MADD, a 15-minute daily exercise regimen led to improvements in locomotor and cardiac function. Although the FI improvement was statistically significant in only L127R mutants **(Fig. 5 B)**, improved cardiac performance was evident in all three mutants in terms of contractility and rhythmicity **(Fig. 7 A-H)**. These functional gains were accompanied by a marked reduction in lipid droplet accumulation as revealed by immunofluorescence imaging **(Fig. 5C-D)** suggesting that physical activity is beneficial in curtailing lipid overload. Exercise was associated with a reduced oxidative burden **(Fig. 5 E-F)** along with elevated expression of mitochondrial remodeling–related genes such as Srl (PGC-1α) and Tfam **(Fig. 5G-I)**. Together, these findings support a framework in which exercise engages adaptive programs linked to mitochondrial remodeling that may help buffer lipid- and ROS-associated mitochondrial dysfunction, rather than directly reversing the underlying genetic defect. These observations are summarized in a working model **(Fig. 7I)**. ETFDH-linked mutations promote lipid accumulation and oxidative stress, which are associated with impaired mitochondrial function, reduced respiration, and decreased ATP synthesis. These metabolic disturbances contribute to progressive contractile dysfunction and are accompanied by activation of cellular stress-response pathways. Consistent with this model, mutant flies exhibited increased expression of AMPK and PGC-1α, together with elevated levels of the mitochondrial remodeling markers Tfam and Opa1, suggesting engagement of an adaptive mitochondrial biogenesis and remodeling program. Increased expression of Sod2 further indicates activation of antioxidant defense mechanisms, whereas upregulation of Pink1 and Parkin is consistent with enhanced mitochondrial quality-control responses triggered by mitochondrial stress. Exercise is proposed to reinforce these adaptive pathways through activation of AMPK-PGC-1α signaling, promoting mitochondrial remodeling and protective stress-response mechanisms that partially counteract oxidative stress, mitochondrial dysfunction, and the physiological consequences of ETFDH deficiency **(Fig. 7I)**

Our study is consistent with previous works in mammalian and *Drosophila* models^78^, where exercise has been shown to activate AMPK signaling, enhance mitochondrial biogenesis, and improve muscle endurance and metabolic flexibility in the context of mitochondrial dysfunction^71^. Memme et al. in 2021 also demonstrated that endurance exercise promotes mitochondrial remodeling and improves muscle oxidative capacity in models of mitochondrial myopathy^79^. Our findings hold significant clinical significance, especially given the limited efficacy of current treatments for MADD, such as riboflavin supplementation and dietary fat restriction^44^. However, clinical translation should be variant-informed and individualized. Late-onset MADD commonly presents with exercise intolerance which improves in many cases after riboflavin therapy^15, 55, 56^. These observations support moderate, supervised physical activity as a complementary strategy once biochemical stabilization is achieved. We infer that moderate aerobic activity can amplify the same mitochondrial biogenesis/oxidative remodeling programs observed in our fly data, provided intensity and duration are carefully tailored to avoid decompensation. In contrast, ETF A/ETF B variants and ETFDH genotypes with null or severely destabilizing alleles are more often associated with severe neonatal or early-onset disease with minimal residual activity, in which exercise is unlikely to be beneficial and may pose rhabdomyolysis risk^16, 80^.

## Conclusion

This study presents an innovative *Drosophila* as a model for investigating the pathogenesis and therapeutic modulation of LSMs, specifically MADD caused by ETFDH mutations in progressive manner. We introduce clinically relevant missense mutations (L127R, S296C, and L399F) into the conserved FAD and ubiquinone-binding domains of the fly Etf-QO gene. We successfully recapitulated hallmark clinical features of MADD, including progressive neuromuscular decline, ectopic lipid accumulation, cardiac dysfunction, and mitochondrial stress. Unlike previous models that rely on acute knockdown or pharmacological inhibition, our CRISPR-engineered lines carry stable, patient-relevant mutations, enabling longitudinal assessment of disease progression and gene-environment interactions. The age-dependent worsening of motor and cardiac phenotypes, coupled with lipid dysregulation and myofibrillar disorganization, mirrors the clinical trajectory of late-onset MADD and provide mechanistic insights into how mitochondrial dysfunction drives tissue degeneration and activates compensatory mitochondrial remodeling, antioxidant defense, and mitochondrial quality-control pathways.

A key translational finding of this work is the demonstration that moderate, daily exercise significantly improves muscle performance and cardiac function in Etf-QO mutants. These benefits were accompanied by reduced lipid accumulation and improved locomotor performance suggesting that exercise promotes mitochondrial remodeling and metabolic reprogramming. Furthermore, the observed reduction in oxidative stress markers highlights the potential of exercise to restore redox balance, positioning it as a promising non-pharmacological intervention for mitochondrial lipid disorders. Our findings not only validate the *Drosophila* model as a powerful platform for dissecting the molecular underpinnings of ETFDH-related myopathies but also open avenues for high-throughput screening of therapeutic compounds and genetic modifiers. This innovative *Drosophila* model demonstrates that even short bouts of moderate exercise can yield significant cardiac benefits, reinforcing its utility for studying gene-environment interactions and non-pharmacological interventions in mitochondrial disease. This model offers a unique opportunity to explore combinatorial interventions that integrate exercise, dietary modulation, and antioxidant therapies.

## Limitations

While exercise improved locomotor and cardiac function and reduced lipid burden, the underlying molecular mechanisms were only partially characterized. Although transcriptional evidence supports activation of AMPK signaling, mitochondrial biogenesis, antioxidant responses, and Pink1-Parkin–associated mitochondrial quality control, direct measurements of mitophagic flux, mitochondrial turnover, and pathway activity were not performed. Our findings demonstrate that a short daily negative geotaxis–based exercise improves LSM phenotypes in the *Drosophila* model; however, direct translation to human therapy is challenging. The 15-minute regimen represents a substantial proportion of the fly’s lifespan and metabolic capacity, whereas human exercise prescriptions must account for differences in body size, energy expenditure, and cardiovascular physiology. Therefore, these results should be interpreted as proof-of-concept for the potential benefits of moderate physical activity rather than a direct recommendation for duration or intensity in patients. Future work should include unbiased transcriptomics and lipidomic to define pathways and lipid species linked with LSM progression and exercise response, which is beyond the scope of this study. Studies in mammalian models and, ultimately, clinical trials are needed to establish optimal exercise parameters for ETFDH-related myopathies. Additional efforts should test combinatorial strategies, including dietary modulation and circadian-based interventions, to enhance translational relevance.

## Availability of data and material

All the data associated with the manuscript has been presented as source data.

## Authors’ information and contribution

Under the GCM guidelines, YG, MD and SB outlined the manuscript, YG and MD initially generated data for this manuscript. SB did nearly all the experiments and wrote the manuscript and generated the figures. GCM edited and revised the manuscript including feedback from all the authors.

## Acknowledgement

We would like to thank Dr. Mona Abdelhamid and Mr. John Watson (Melkani Lab) for their help in isolating RNA from some of the *Drosophila* thorax samples and help with the fluorescence microscopy respectively.

