## Supplementary material for "Innovative CRISPR Etf-QO Mutant Knock-in Model Mimics MADD-associated Lipid Storage Myopathy and Reveal Therapeutic Potential of Exercise Intervention": NA

### ***Supplementary Information***

#### ***Modeling Late-Onset Multiple Acyl-CoA Dehydrogenase Deficiency in Drosophila Identifies Exercise-Driven Rescue of Lipid Storage Myopathy***

Sachin Budhathoki [a], Yiming Guo [a], Mary Doamekpor [a], and Girish Melkani [a,b,*]

[a]Department of Pathology, Division of Molecular and Cellular Pathology, Heersink School of Medicine, Heersink School of Medicine, The University of Alabama at Birmingham, AL 35294, USA. [b]UAB Nathan Shock Center, The University of Alabama at Birmingham, AL 35294, USA

[*]Correspondence Department of Pathology, Division of Molecular and Cellular Pathology, School of Medicine, University of Alabama at Birmingham, AL 35294, USA. Tel.: 1-205-996-0591; Fax: 1-205-934-7447;

Number of figures: 5

Fig. SI1

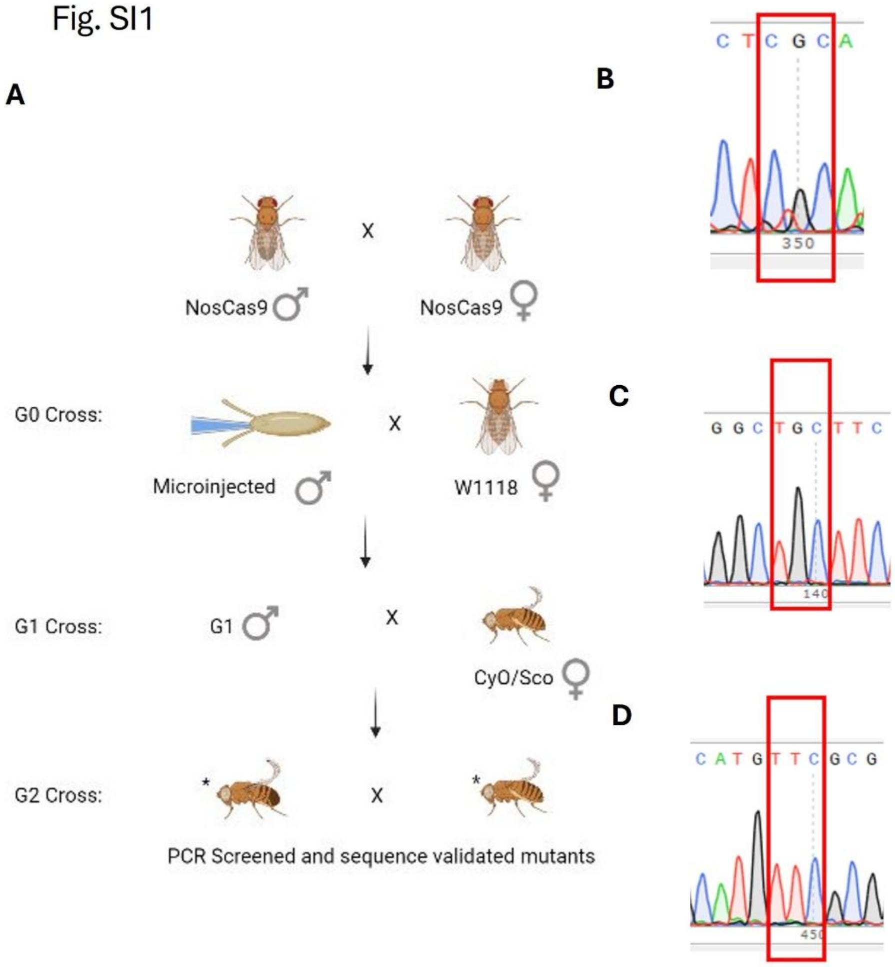


**Figure SI1.** Generation and validation of CRISPR knock-in Drosophila models carrying ETFQO point mutations associated with Multiple Acyl-CoA Dehydrogenase Deficiency (MADD). **(A)** Schematic representation of the CRISPR/Cas9-based workflow for introducing targeted point mutations in the *Etf-QO* gene. sgRNA-target plasmids and donor templates containing the desired mutations were microinjected into nos-Cas9 embryos (G0), followed by successive genetic crosses using balancer chromosomes to establish stable mutant lines. **(B–D)** Representative Sanger sequencing chromatograms confirming the presence of specific point mutations: L127R **(B)**, S296C **(C)**, and L399F **(D)**, located within conserved FAD and ubiquinone (UQ) binding domains of ETFQO.

### Fig. SI2

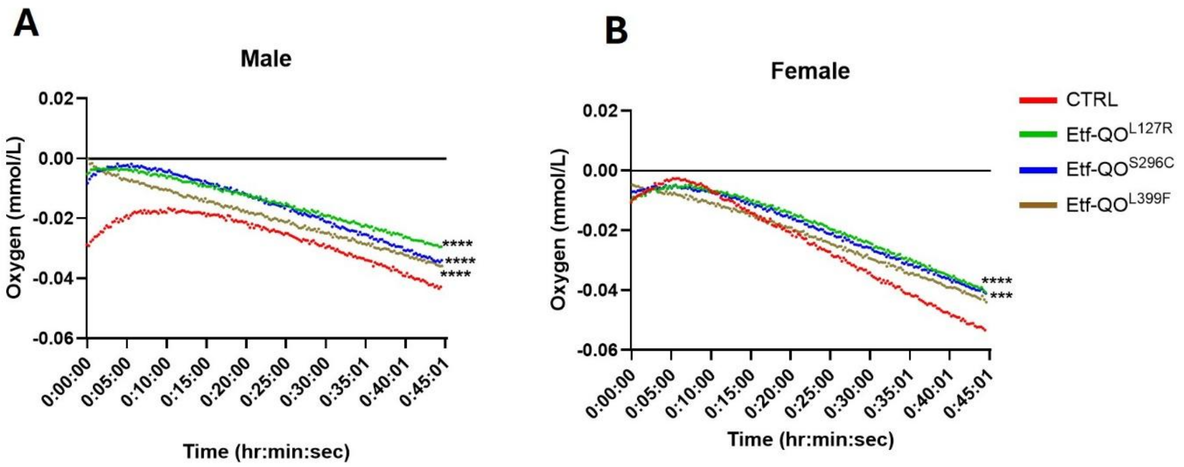


**Figure SI2. Assessment of the effect of ETF-QO mutantion in mitochondrial respiration.** Oxygen concentration profiles over time in exercise-conditioned male (A) and female (B) flies from control (CTRL) and ETF-QO mutant genotypes (**L127R**, **S296C**, **L399F**) measured using the MicroResp™ respirometry system (Loligo Systems). Each curve represents mean oxygen concentration (mmol/L) recorded at 1-second intervals for 45 minutes under standardized conditions. Mutants exhibited lower oxygen consumption rates compared to controls, indicating impaired respiratory capacity. Blank chambers served as negative controls. Data was analyzed using MicroResp software. Sample data corrected by subtracting the blank mean oxygen value from individual oxygen value across time. n = 5 flies per group. One-way ANOVA with Dunnett's multiple comparisons test, **p<0.01, ***p<0.001, ***p<0.0001

Fig. SI3

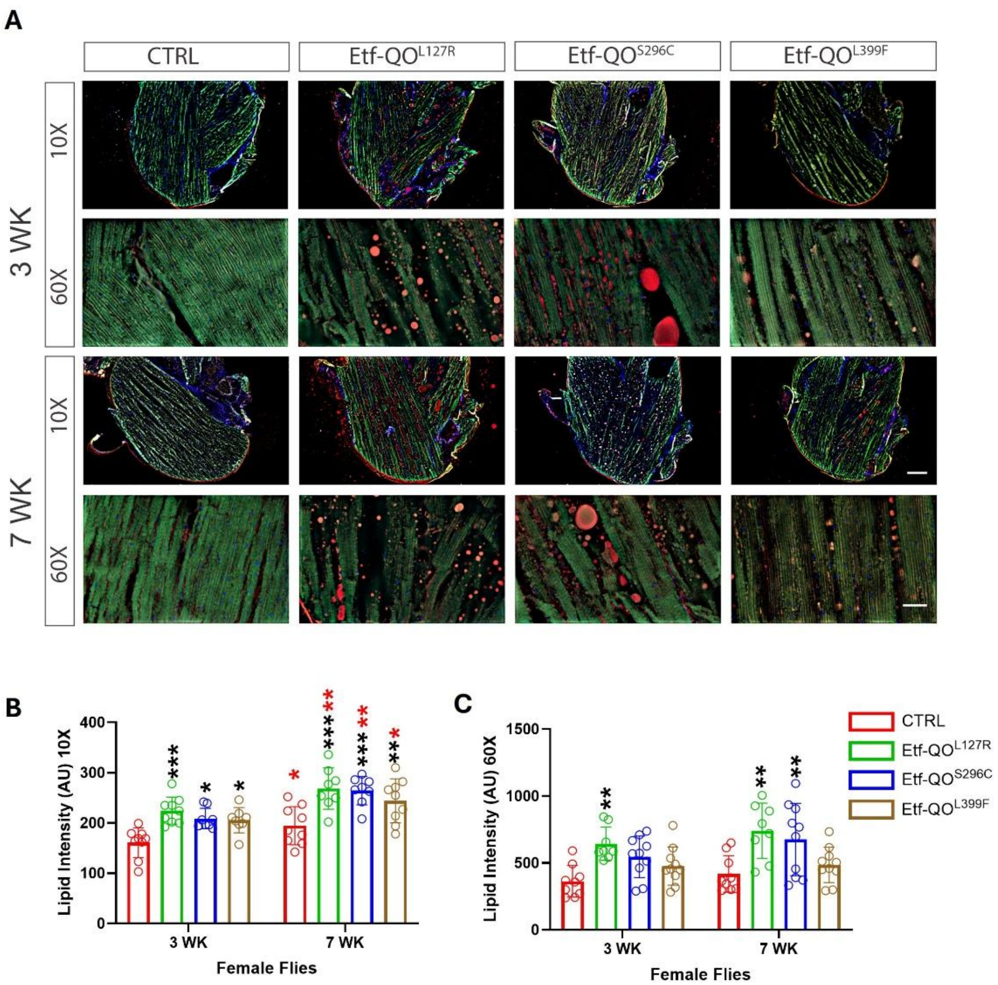


**Figure SI3.** Representative sagittal cryosections of thoracic muscles from control and Etf-QO mutant female flies at mid age (3 weeks) and old age (7 weeks) age, imaged at 10X and 60X magnification **(A)**. Sections were stained with phalloidin (green) to visualize actin cytoskeleton and DAPI (blue) for nuclei. Lipid droplets appear as red puncta. Mutants exhibit markedly higher lipid accumulation compared to controls, with severity slightly increasing at 7 weeks **(B)**. Quantification of lipid intensity at 10X **(B)** and 60X **(C)** magnification confirms significant elevation in lipid deposition in mutants relative to controls across both ages. (n=10–20 images, 5 females/group). Two-way ANOVA with Tukey's multiple comparisons test, $**p<0.01$, $***p<0.001$. Scale bar:100 μm(10X), 20 μm(60X).

Fig. SI4

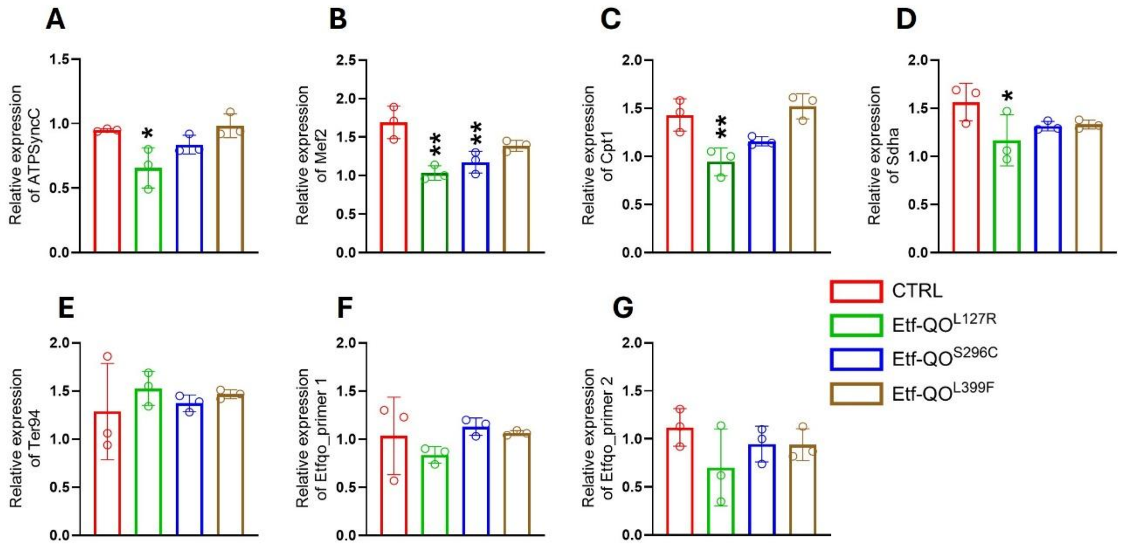


**Figure SI4. Etf-QO mutations alter metabolic and muscle-maintenance transcripts.** Relative mRNA expression levels of ATP synthase subunit C (A), Mef2 (B), Cpt1 (C), SdhA (D), Ter94 (E), and Etf-QO measured using two independent primer pairs (F-G). L127R exhibited reduced expression of AtpsynC, Mef2, Cpt1, and SdhA, consistent with impaired ATP production, fatty-acid utilization, mitochondrial metabolism, and muscle maintenance. Ter94 showed minimal genotype-dependent changes, whereas Etf-QO transcript abundance remained broadly comparable among alleles using both primer sets, indicating that the observed phenotypes arise primarily from functional consequences of the missense mutations rather than altered transcript abundance. One-way ANOVA with Dunnett's multiple-comparisons test; *$p < 0.05$,** $p < 0.01$, ***$p < 0.001$

Fig. SI5

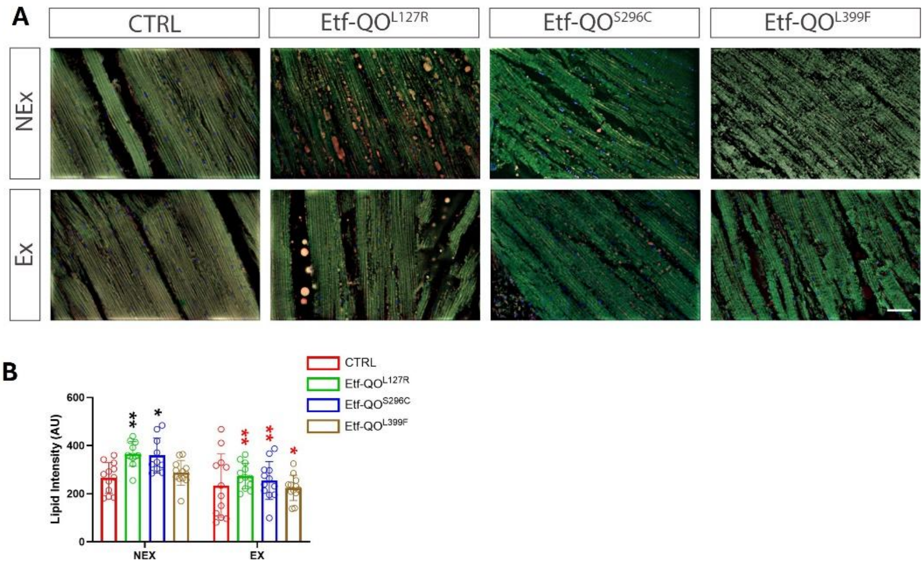


**Figure SI5**. **Effect of moderate exercise on lipid accumulation in indirect flight muscles of ETF-QO mutants**. Representative fluorescent images of thoracic indirect flight muscles from control (CTRL) and ETF-QO mutant flies under non-exercise (NEX) and exercise (EX) conditions **(A)**. Muscles were stained with phalloidin (green) to visualize actin filaments; lipid droplets appear as red puncta. Regular exercise (15 min/day for 2.5 weeks) visibly reduced lipid deposition in all mutant genotypes compared to sedentary cohorts. Scale bar: 100 µm. Quantification of lipid object count **(B)** and lipid object area **(C)** from IFM images. (n=10–20 images, 5 males/group). Two-way ANOVA with Tukey's multiple comparisons test, **p<0.01, ***p<0.001. Black and red asterisks denote comparison across genotypes and exercise conditions, respectively. Scale bar: 20 µm

### Fig. SI6

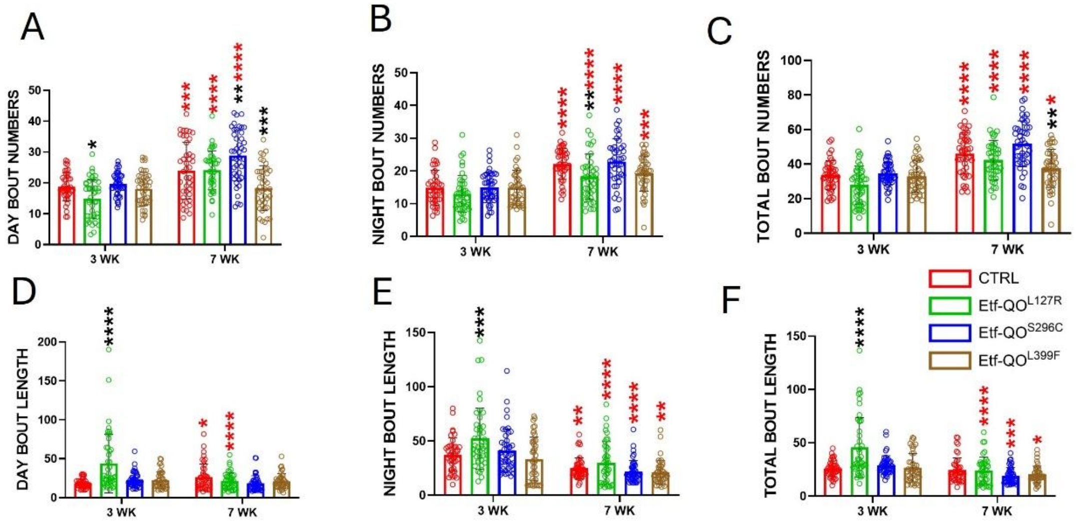


**Figure SI6. Etf-QO mutations alter sleep architecture.** Daytime (A), nighttime (B), and total bout number (C) demonstrate genotype-dependent alterations in sleep organization. Daytime (D), nighttime (E), and total bout length (F) further indicate differences in sleep consolidation among mutant backgrounds. Across most measurements, L127R exhibited the strongest phenotype, S296C showed intermediate effects, and L399F displayed comparatively mild abnormalities. Data are presented as mean ± SD. Statistical significance was determined using one-way or two-way ANOVA with appropriate post hoc testing. All raw data are provided in the Source Data file.
